# Alternative *REST* Splicing Underappreciated

**DOI:** 10.1101/119552

**Authors:** Guo-Lin Chen, Gregory M. Miller

## Abstract

As a major orchestrator of the cellular epigenome, the repressor element-1 silencing transcription factor (REST) can either repress or activate thousands of genes depending on cellular context, suggesting a highly context-dependent REST function tuned by environmental cues. While *REST* shows cell-type non-selective active transcription^1^, an N-terminal REST4 isoform caused by alternative splicing – inclusion of an extra exon (N_3c_) which introduces a premature stop codon – has been implicated in neurogenesis and tumorigenesis^2-5^. Recently, in line with established epigenetic regulation of pre-mRNA splicing^6,7^, we demonstrated that *REST* undergoes extensive, context-dependent alternative splicing which results in the formation of a large number of mRNA variants predictive of multiple protein isoforms^8^. Supported by that immunoblotting/-staining with different anti-REST antibodies yield inconsistent results, alternative splicing allows production of various structurally and functionally different REST protein isoforms in response to shifting physiological requirements, providing a reasonable explanation for the diverse, highly context-dependent REST function. However, REST isoforms might be differentially assayed or manipulated, leading to data misinterpretation and controversial findings. For example, in contrast to the proposed neurotoxicity of elevated nuclear REST in ischemia^9^ and Huntington’s disease^10,11^, Lu et al. recently reported decreased nuclear REST in Alzheimer’s disease and neuroprotection of REST in ageing brain^12^. Unfortunately, alternative *REST* splicing was largely neglected by Lu et al., making it necessary for a reevaluation of their findings.

---

As shown in Fig.1a, human *REST* gene boundary is now doubled by an alternative last exon (E_5_) which is mutually exclusive to E_4_. While numerous novel alternative exons and 5’/3’ ends were identified, the 3 constitutive exons (E_2_, E_3_ and E_4_) comprising the open reading frame (ORF) of *REST* can be skipped partially or completely, alone or in combination, producing at least 45 mRNA variants predictive of multiple protein isoforms (Fig.1b)^8^. For example, REST4 – which was first described in rat as a group of REST isoforms^4^, is predicted by multiple mRNA variants (e.g. JX896958, JX896971 and JX896983) with E_3_ followed by variable exons introducing a premature stop codon, suggesting that like the case in rat, human REST4 is also a group of isoforms, but not a single mRNA/protein isoform. Meanwhile, REST1 – another N-terminal isoform is predicted by multiple mRNA variants lacking E_3_. Notably, for E_2_-skipped variants (e.g. XM_005265760 and JX896960) missing the conventional start codon, an in-frame AUG in E_3_ may initiate translation of a C-terminal REST^C^ isoform (XP_005265817), which was recently described in *Rest* conditional knockout (cKO) mice^13^. In addition, some mRNA variants (e.g. JX896978 and KC117266) with partial E_2_ skipping are predictive of proteins missing variable regions of REST. Moreover, it was recently demonstrated that mRNAs with short ORF but previously annotated as noncoding RNAs can encode tiny peptides^14-16^, such might be the case for numerous REST variants (e.g. JX896962, JX896965, and JX896967). Taken together, REST protein isoforms caused by alternative splicing are much more complex than we expected.

Due to the existence of multiple REST mRNA/protein isoforms, it can be inferred that assay of REST expression by different primers (or probes) and antibodies may target different REST isoforms, while manipulation of REST expression by cKO or RNAi may be effective for specific but not all REST variants. In other words, REST isoforms might be differentially assayed or manipulated in different studies, leading to inconsistent results and data misinterpretation. In support of this notion, immunostaining of multiple cell lines with two widely used antibodies – sc-25398 and ab21635 raised against N- and C-terminus of REST, respectively – produced inconsistent results in terms of REST subcellular distribution and its co-localization with microtubule^11,17^ (Fig.2), while immunoblotting (i.e. Western blotting or WB) with the two antibodies yielded different profiles of immunoreactive (IR) bands (Fig.3), such is the case for some other commercial anti-REST antibodies as described by manufacturer’s manual. Unfortunately, despite the mRNA evidence, not all REST protein isoforms have been experimentally verified and they are usually observed as unexpected sizes due to post-translational modifications, making it challenging to determine whether an unknown IR band is non-specific or a REST isoform. For example, REST4 and REST^C^ are predicted as 37 and 86 kD but observed as 53 and 130 kD, respectively^3,13^, while the full-length REST has been reported as variable sizes ranging from 120 to 200 kD^13,18,19^. So, even if detectable by WB, specific REST isoforms might be simply considered as non-specific and excluded from being presented in publication, such may explain why REST^C^ was not reported until recently.

**Fig. 1.**
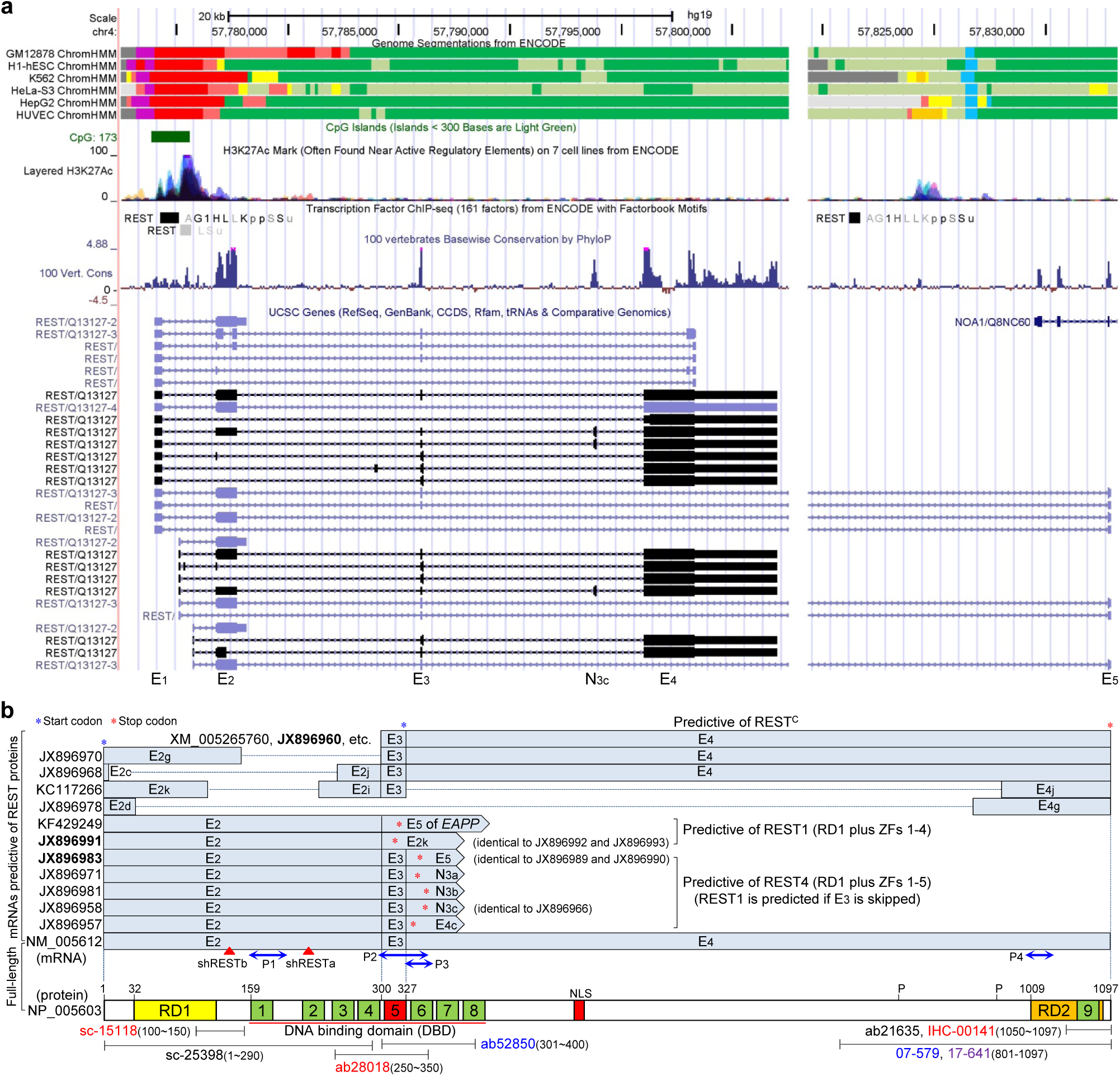
Bioinformatics at human *REST* locus (a) and predicted REST protein isoforms derived from alternative splicing (b). Informative tracks were retrieved from the UCSC Genome Browser (http://genome.ucsc.edu). *REST* gene boundary is largely expanded by the alternate last exon (E_5_) which partially overlaps in opposite direction with exon 5 of *NOA1*. *REST* promoter harbors a CpG island and exhibits cell-independent active transcription as indicated by information of the chromatin state segmentation and H3K27Ac tracks. Predicted open reading frames (ORFs) of the full-length and alternatively spliced REST mRNAs were briefly shown by indicating the start (blue star) and stop (red star) codons, while major domains (RD1/RD2 – repression domain 1 and 2; NLS – nuclear localization signal; and zinc fingers 1-9) of the full-length REST protein were proportionally illustrated in parallel to their coding sequences. Locations of the mRNA and protein fragments targeted by real-time PCR primer sets (P1-P4), RNAi (shRESTa and shRESTb) and antibodies mentioned in the text were indicated. Note that: 1) there are multiple alternative transcription start sites (only 3 well-documented were shown) and not all RNA variants are listed; 2) ChIP-seq identified 3 RE-1 sites across *REST* locus; and 3) the internal region of E_4_ is non-conservative as indicated by the “100 Vertebrate Conservation” track, which supports our finding that partial skipping of E_4_ is common.

**Fig. 2.**
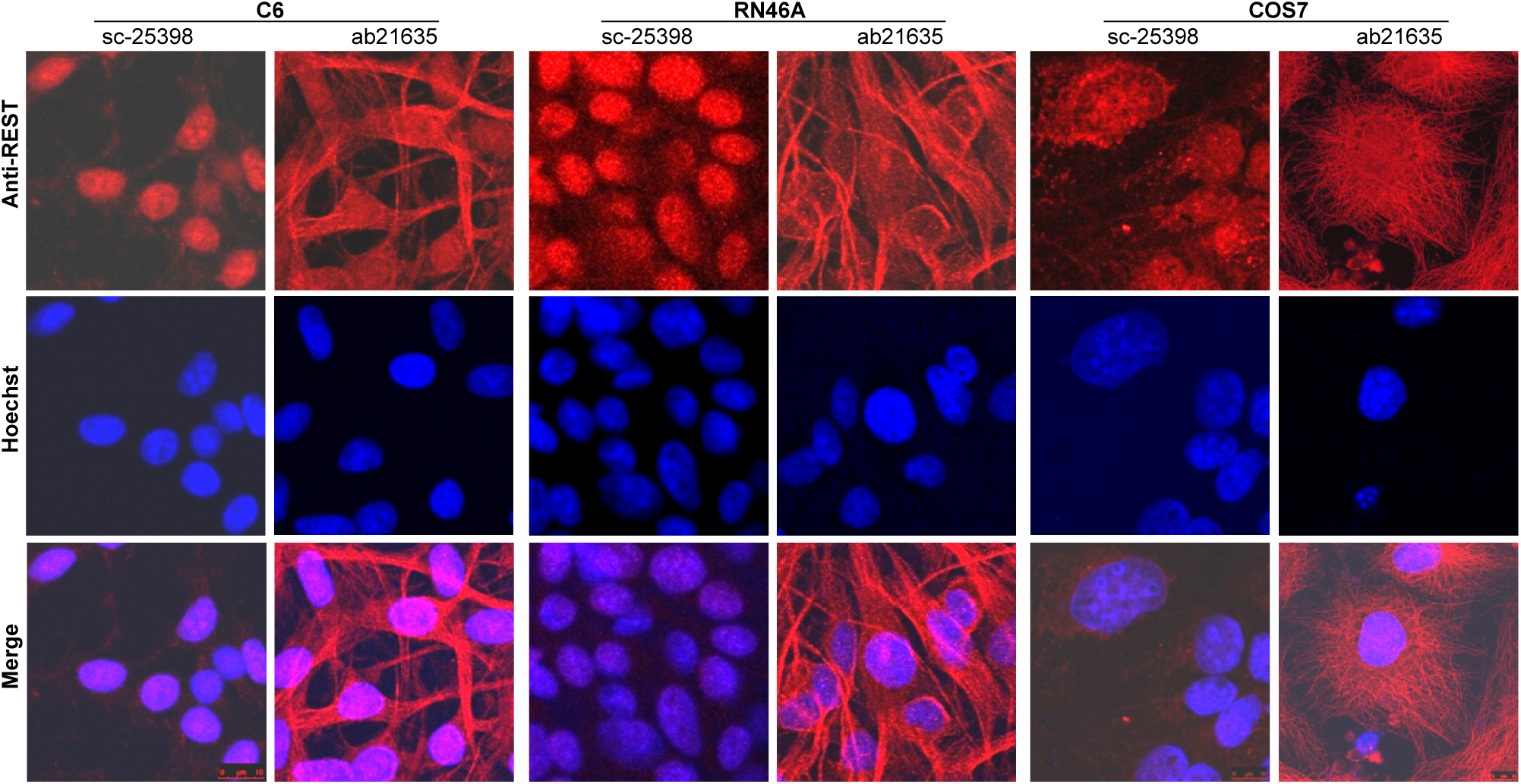
Immunofluorescence analysis of REST subcellular localization in different cells with two different antibodies. ICC assays were performed with two anti-REST antibodies (sc-25398 and ab21635, against N- and C-terminus of REST, respectively) for C6, RN46A, and COS7 cells. For each cell line, cells cultured in two wells of 24-well plate were processed for ICC analysis with sc-25398 and ab21635, respectively, with all the experimental conditions (e.g. passage number and density of the cells, culture condition and ICC procedure) were kept the same for the two antibodies. It is obvious that regardless of the cell-types, ICC with sc-25398 yielded predominant localization of REST in nucleus, whereas ICC with ab21635 indicated predominant co-localization of REST with microtubule (or cytoskeleton), suggesting that REST isoforms with different subcellular localization might be differentially recognized by different antibodies. Dr. Qi Ma at SUNY Upstate Medical University provided technical support for the ICC analysis.

**Fig. 3.**
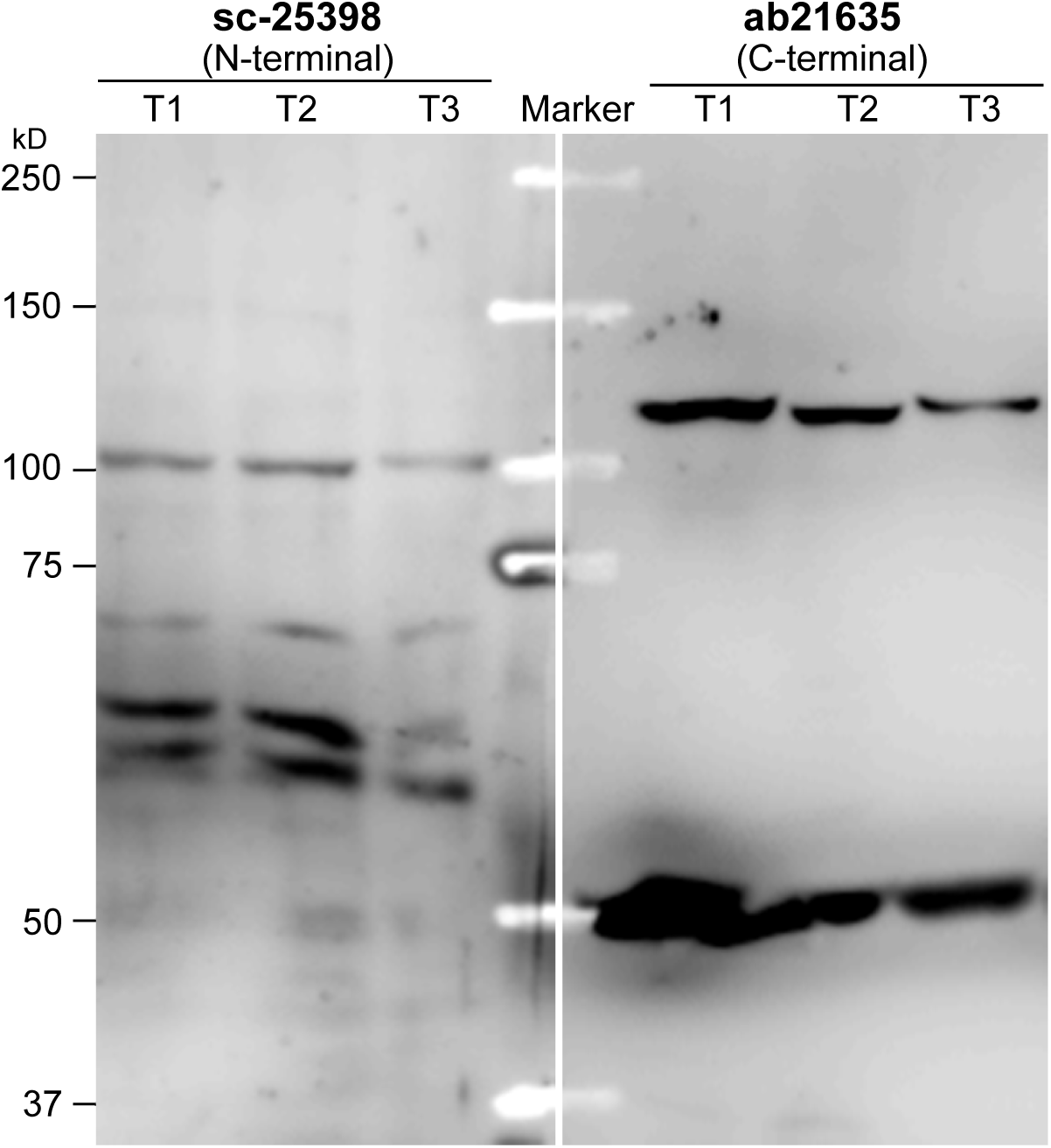
Western blotting analysis of REST expression in HEK-293T cells with two different antibodies. Two aliquots (25μg for each) of 3 different HEK293T protein samples (T1, T2 and T3), which were isolated simultaneously with RNA and DNA by TRIzol^®^ Reagent (Invitrogen), along with a Kaleidoscope(tm) marker (Bio-Rad) in between, were loaded on a 7.5% PAGE-SDS gel, followed by electrophoresis and electrotranslocation onto an Immun-Blot PVDF membrane (Bio-Rad), which was then cut into two halves for incubation with sc-25398 (1:250) and ab21635 (1:500), respectively, and subsequent incubation with a goat anti-rabbit IgG (Sigma-Aldrich, 1:2500). Immunoreactive signals were detected using the VisiGlo^TM^ Select HRP Chemiluminescent Substrate Kit (Amresco) with an ECL-based LAS-3000 image system (Fujifilm). Note that the two antibodies yielded totally different profiles of IR bands.

In their paper describing altered nuclear REST in ageing and AD brain, Lu et al. claimed that REST4 mRNA (N_3c_) level in brain tissues comprised only 0.1-0.5% of *REST* mRNA^12^, while splice variants caused by ΔE_2_, ΔE_3_ and ΔE_4_ (or inclusion of E_5_) exhibit neuronal expression, of which ΔE_3_ eliminates a motif critical for nuclear targeting^20,21^ and therefore affects nuclear REST^22^) were not mentioned. It can be simply inferred that if only the full-length REST mRNA exists, all segments of it should share the same level of expression; however, in accordance with our above-mentioned notion of inconsistent results yielded by different primers, qRT-PCR data in Lu et al. indicated that 4 primer sets (P1-P4) targeting different exons of *REST* yielded strikingly different mRNA expression change. Notably, patterns of this primer-dependent result varied across the aged groups. For instance, P2 assay showed the highest and lowest fold change for the 95-yr and >95-yr group, respectively, while some assays for ageing groups (e.g. P1/P4 for 71-yr, P1/P3/P4 for 95-yr, and P2 for >95-yr) showed similar levels of mRNA expression with the 25-yr group, suggesting that systematic error made minor contribution to this inconsistence, which however can be explained by our previous finding of individual variation in alternative *REST* splicing^8^. So, qRT-PCR data presented by Lu et al. actually provided strong evidence for alternative *REST* splicing, which however was not interpreted by the authors. In addition, unlike Northern blotting which gives size information for observed mRNAs, qRT-PCR measures abundance of a specific amplicon (i.e. a segment of mRNA) which might be shared by multiple mRNA variants, such that qRT-PCR data represents expression of all splice variant(s) yielding the same amplicon, but not merely the full-length *REST* mRNA. Hence, without any evidence of Northern blotting, it’s difficult to interpret the full-length *REST* mRNA expression with the primer-dependent qRT-PCR data in Lu et al. Meanwhile, Lu et al. performed a series of experiments (e.g. RNAi, ChIP-seq and oxidative stress) with the SH-SY5Y cell line which reportedly expresses abundant REST4 mRNA (N_3c_) and protein^5,8,23^; however, REST4 expression in SH-SY5Y cells was not mentioned in the paper.

Lu et al. assayed REST protein expression level and subcellular distribution by immunoblotting/-staining with a total of 6 different antibodies, of which 2 (07-579 and ab52850) and 3 (ab28018, sc-15118 and IHC-00141) were used for Western and immunohistochemistry (IHC), respectively. As mentioned above, due to the existence of multiple *REST* protein isoforms, different antibodies may yield different Western/IHC results, while Western with a specific antibody may yield multiple IR bands which represent different REST isoforms sharing the same epitope. So, comparison of the Western/IHC results between different antibodies may hint about the existence of multiple REST protein isoforms; however, Lu et al. did not show any comparison of the Western/IHC results between different antibodies, and all the presented blots were cropped with only the band of interest (presumably represents the full-length REST) observable, making it impossible to evaluate the existence of multiple REST, even for the established REST4 in SH-SY5Y cells. Although Lu et al. performed immunostaining to test specificity of one IHC antibody (IHC-00141), it cannot exclude the existence of multiple REST isoforms, because isoforms sharing the same epitope can all bind to the same antibody, and this binding can be eliminated by the same blocking peptide.

Notably, it was also not disclosed how the 3 IHC antibodies were assigned to samples of different groups in Lu et al., giving rise to the concern that nuclear REST differences between the experimental groups might be “artificially generated” by biased usage of the antibodies for different samples. For example, comparison of nuclear REST between young (n=11), aged (n=77), AD (n=72), and MCI (n=11) groups (Fig.1e-Imaging in Lu et al.) was presumably based on staining of the samples with 3 different antibodies but not a single antibody, otherwise the remaining 2 antibodies must have been respectively used for another two sets of samples, which however were totally not mentioned in the paper. So, without considering differences between the antibodies and disclosing usage of the antibodies, the employment of multiple antibodies for IHC did not strengthen findings of Lu et al., but instead introduced an extra confounding variable which made the findings questionable.

In response to our doubt about the usage of the antibodies, *Nature* published an addendum on 16 November 2016^24^. Specifically, as shown in Table 1, several occasions of IHC experiments – which had not been previously mentioned – were added to the article, making that each antibody was seemingly used on an independent occasion of IHC experiment. While this addendum provided information essential for follow-up research, it raised other concerns: **1)** it’s unknown why the added experiments had not been previously mentioned in the article, making it doubtful when these experiments (presumably not peer-reviewed) were performed; **2)** according to the addendum, all the IHC data presented in Lu et al. were obtained with the antibody IHC-00141, while the other two IHC antibodies (and an additional antibody ab202962) yielded similar results (listed as “data not shown”). It goes without saying that similar results of multiple antibodies will make the data more solid and convincing than result of a single antibody; however, it’s strange that only the result of IHC-00141 was presented in Lu et al., while results of the other two antibodies were not described even though all the 3 antibodies were mentioned in the original article ["Immunofluorescence microscopy using three different antibodies against the amino- or carboxy-terminal domains of REST showed a striking induction of REST in the nucleus of ageing neurons in the PFC and hippocampus (Fig. 1d, e and Extended Data Fig. 1)”]; **3)** it was claimed in the addendum that 7 cases from the 171 cases labeled with IHC-00141 were labeled in separate slides with ab28018 and that “similar patterns of REST immunoreactivity were observed with increased nuclear REST in neurons in aged versus young adult cases and reduced nuclear REST in AD”, but the sample size (7 cases comprising at least 3 experimental groups, i.e. n<3 for 1-2 groups) is extremely too small to perform statistics; and **4)** like the case for IHC, two antibodies (07-579 and ab52850) were previously listed for WB without disclosure of their usage for each independent experiment; however, based on the addendum, 07-579 was used to generate all the Western data presented in Lu et al, while neither the usage nor the result information was disclosed for ab52850, making it questionable for the rationale of listing ab52850 in the paper.

**Table 1.**
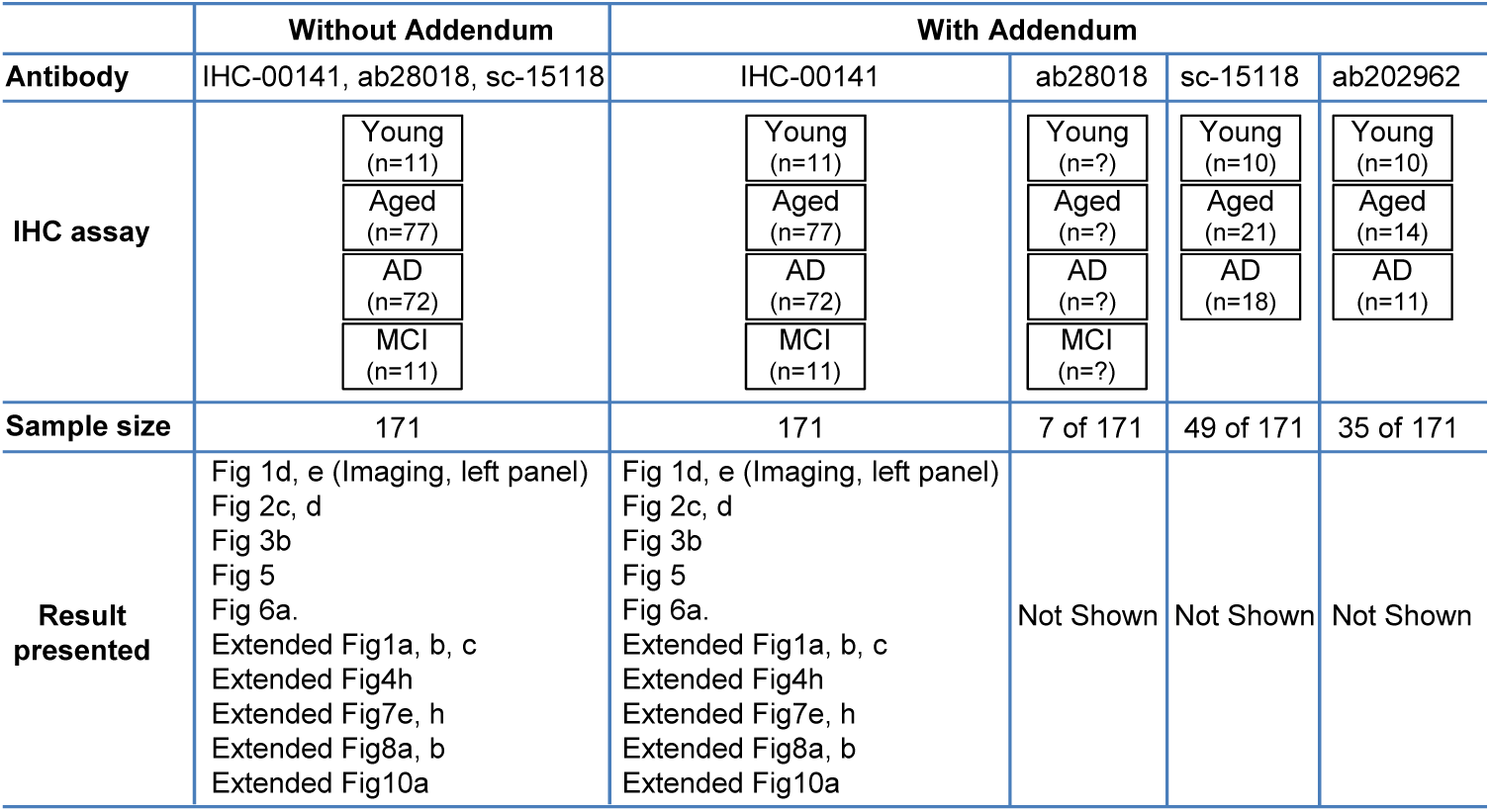
A summary of IHC assay mentioned in Lu et al. with and without the Addendum.

|  | Without Addendum | With Addendum |  |  |  |
| --- | --- | --- | --- | --- | --- |
| Antibody | IHC-00141, ab28018, sc-15118 | IHC-00141 | ab28018 | sc-15118 | ab202962 |
| IHC assay | Young<br>(n=11) | Young<br>(n=11) | Young<br>(n=?) | Young<br>(n=10) | Young<br>(n=10) |
|  | Aged<br>(n=77) | Aged<br>(n=77) | Aged<br>(n=?) | Aged<br>(n=21) | Aged<br>(n=14) |
|  | AD<br>(n=72) | AD<br>(n=72) | AD<br>(n=?) | AD<br>(n=18) | AD<br>(n=11) |
|  | MCI<br>(n=11) | MCI<br>(n=11) | MCI<br>(n=?) |  |  |
| Sample size | 171 | 171 | 7 of 171 | 49 of 171 | 35 of 171 |
| Result presented | Fig 1d, e (Imaging, left panel)<br>Fig 2c, d<br>Fig 3b<br>Fig 5<br>Fig 6a.<br>Extended Fig1a, b, c<br>Extended Fig4h<br>Extended Fig7e, h<br>Extended Fig8a, b<br>Extended Fig10a | Fig 1d, e (Imaging, left panel)<br>Fig 2c, d<br>Fig 3b<br>Fig 5<br>Fig 6a.<br>Extended Fig1a, b, c<br>Extended Fig4h<br>Extended Fig7e, h<br>Extended Fig8a, b<br>Extended Fig10a | Not Shown | Not Shown | Not Shown |

Even if a fixed antibody was used throughout the study, expression of multiple REST isoforms caused by alternative splicing may lead to data misinterpretation. For example, the N-terminal REST4, whose expression in SH-SY5Y was ignored by Lu et al., competes with the full-length REST to occupy RE-1 sites, such that it inevitably affects interpretation of REST target genes with the ChIP-seq data. Also, REST isoforms sharing the same epitope can be indiscriminately labeled by a specific antibody, and in comparison with the full-length REST, truncated isoforms might be more easily accessible to the antibody due to less complexity of protein folding and 3-dimentional structure which potentially masks the epitope. As mentioned above, test of antibody specificity by immunostaining does not simple help to exclude the existence of multiple REST isoforms sharing the same epitope, whose binding to the antibody can be eliminated by the same blocking peptide. So, the IHC results could not address which specific REST isoform(s) contributed to differences in nuclear REST between the experimental groups; however, with only the full-length REST having been considered, such differences were attributed to the full-length REST in Lu et al.

Taken together, Lu et al. neglected previously documented REST isoforms which presumably confound experimental results and lead to data interpretation, while an extra confounding variable was introduced in that study by employing multiple antibodies to assay REST protein expression, making that findings of Lu et al. are questionable and require a re-evaluation.

## Acknowledgments

This work was supported by NIH grants DA030177 (GMM). We thank Dr. Qi Ma at SUNY Upstate Medical University for technical support for the ICC analysis.

### Conflict of interest

Both of the authors do not have any financial disclosures to report.

## Figure Legends

**Fig.1b. Bioinformatics at human *REST* locus (a) and predicted REST protein isoforms derived from alternative splicing (b)**. Related tracks were retrieved from the UCSC Genome Browser (http://genome.ucsc.edu/cgi-bin/hgGateway). *REST* gene boundary is more than doubled by an alternate last exon (E_5_) which partially overlaps in opposite direction with exon 5 of *NOA1*. *REST* promoter harbors a CpG island and exhibits cell-independent active transcription as indicated by the chromatin state segmentation and H3K27Ac tracks. Predicted open reading frames (ORFs) of the full-length and alternatively spliced REST mRNAs were briefly shown by indicating the start (blue star) and stop (red star) codons, while major domains (RD1, RD2 – repression domain 1 and 2; NLS – nuclear localization signal; and zinc fingers 1-9) of the full-length REST protein were illustrated in parallel to their coding sequences. Splice variants expressed in multiple tissues or cell lines were bolded. Locations of the mRNA and protein fragments targeted by real-time PCR primer sets (P1-P4), RNAi (shRESTa and shRESTb), and antibodies mentioned in the text were indicated. Note that only the conventional promoter is shown, and that the internal region of E_4_ is unconserved as indicated by the “100 Vertebrate Conservation” track, which supports our finding that partial skipping of E_4_ is common^8^.

**Fig.2. Immunofluorescence analysis of REST subcellular localization in different cells with two different antibodies.** ICC were performed with two anti-REST sc-25398 (Santa Cruz) and ab21635 (Abcam) – which are respectively against N- and C-terminus of REST – for C6, RN46A, and COS7 cells. For each cell line, two wells of cells under the same experimental conditions were stained with sc-25398 and ab21635, respectively. Briefly, cells cultured on poly-D-lysine coated coverslips were fixed with 4% paraformaldehyde, permeabilized with 0.3% Triton X-100, and incubated with sc-25398 (1:100) or ab21635 (1:200), followed by incubation with a goat anti-rabbit secondary antibody conjugated with Alexa Fluor^®^ 568 (1:500, Invitrogen). Nuclei were stained with Hoechst-33342 (Thermo scientific) and cells were mounted on glass slides. Confocal microscopy was performed using a Leica TCS SP5 Spectral Confocal Microscope. For each cell line, all experimental conditions were kept the same for the two antibodies. Regardless of the cell-types, ICC with sc-25398 yielded predominant localization of REST in nucleus, whereas ICC with ab21635 indicated predominant co-localization of REST with microtubule (or cytoskeleton), suggesting that REST isoforms with different subcellular localization might be differentially recognized by different antibodies.

**Fig.3. Western blotting analysis of REST expression in HEK-293T cells with two different antibodies.** Two aliquots (25μg for each) of 3 different HEK293T protein samples (T1, T2 and T3), which were isolated simultaneously with RNA and DNA by TRIzol^®^ Reagent (Invitrogen), along with a Kaleidoscope^TM^ marker (Bio-Rad) in between, were loaded on a 7.5% PAGE-SDS gel, followed by electrophoresis and electrotranslocation onto an Immun-Blot PVDF membrane (Bio-Rad), which was then cut into two halves for incubation with sc-25398 (1:250) and ab21635 (1:500), respectively, and subsequent incubation with a goat anti-rabbit IgG (Sigma-Aldrich, 1:2500). Immunoreactive signals were detected using the VisiGlo^TM^ Select HRP Chemiluminescent Substrate Kit (Amresco) with an ECL-based LAS-3000 image system (Fujifilm). Note that the two antibodies yielded totally different profiles of IR bands.

## References

1 Kojima, T., Murai, K., Naruse, Y., Takahashi, N. & Mori, N. Cell-type non-selective transcription of mouse and human genes encoding neural-restrictive silencer factor. Molecular Brain Research 90, 174–186, doi:10.1016/s0169-328x(01)00107-3 (2001).

2 Raj, B. et al. Cross-Regulation between an Alternative Splicing Activator and a Transcription Repressor Controls Neurogenesis. Molecular Cell 43, 843–850, doi:10.1016/j.molcel.2011.08.014 (2011).

3 Lee, J. H., Chai, Y. G. & Hersh, L. B. Expression patterns of mouse repressor element-1 silencing transcription factor 4 (REST4) and its possible function in neuroblastoma. Journal of Molecular Neuroscience 15, 205–214, doi:10.1385/jmn:15:3:205 (2000).

4 Palm, K., Belluardo, N., Metsis, M. & Timmusk, T. Neuronal expression of zinc finger transcription factor REST/NRSF/XRB gene. Journal of Neuroscience 18, 1280–1296 (1998).

5 Palm, K., Metsis, M. & Timmusk, T. Neuron-specific splicing of zinc finger transcription factor REST/NRSF/XBR is frequent in neuroblastomas and conserved in human, mouse and rat. Molecular Brain Research 72, 30–39, doi:10.1016/s0169-328x(99)00196-5 (1999).

6 Luco, R. F., Allo, M., Schor, I. E., Kornblihtt, A. R. & Misteli, T. Epigenetics in Alternative PremRNA Splicing. Cell 144, 16–26, doi:10.1016/j.cell.2010.11.056 (2011).

7 Alló, M. et al. Chromatin and Alternative Splicing. Cold Spring Harbor Symposia on Quantitative Biology 75, 103–111, doi:10.1101/sqb.2010.75.023 (2010).

8 Chen, G.-L. & Miller, G. M. Extensive alternative splicing of the repressor element silencing transcription factor linked to cancer. PLoS ONE 8, e62217, doi:10.1371/journal.pone.0062217 (2013).

9 Noh, K.-M. et al. Repressor element-1 silencing transcription factor (REST)-dependent epigenetic remodeling is critical to ischemia-induced neuronal death. Proceedings of the National Academy of Sciences 109, E962–E971, doi:10.1073/pnas.1121568109 (2012).

10 Zuccato, C. et al. Huntingtin interacts with REST/NRSF to modulate the transcription of NRSE-controlled neuronal genes. Nature Genetics 35, 76–83, doi:10.1038/ng1219 (2003).

11 Buckley, N. J., Johnson, R., Zuccato, C., Bithell, A. & Cattaneo, E. The role of REST in transcriptional and epigenetic dysregulation in Huntington’s disease. Neurobiology of Disease 39, 28–39, doi:10.1016/j.nbd.2010.02.003 (2010).

12 Lu, T. et al. REST and stress resistance in ageing and Alzheimer’s disease. Nature 507, 448–454, doi:10.1038/nature13163 (2014).

13 Nechiporuk, T. et al. The REST remodeling complex protects genomic integrity during embryonic neurogenesis. eLife 5, doi:10.7554/eLife.09584 (2016).

14 Magny, E. G. et al. Conserved Regulation of Cardiac Calcium Uptake by Peptides Encoded in Small Open Reading Frames. Science 341, 1116–1120, doi:10.1126/science.1238802 (2013).

15 Nelson, B. R. et al. A peptide encoded by a transcript annotated as long noncoding RNA enhances SERCA activity in muscle. Science 351, 271–275, doi:10.1126/science.aad4076 (2016).

16 Olexiouk, V. et al. sORFs.org: a repository of small ORFs identified by ribosome profiling. Nucleic Acids Research, doi:10.1093/nar/gkv1175 (2015).

17 Shimojo, M. Huntingtin Regulates RE1-silencing Transcription Factor/Neuron-restrictive Silencer Factor (REST/NRSF) Nuclear Trafficking Indirectly through a Complex with REST/NRSF-interacting LIM Domain Protein (RILP) and Dynactin p150(Glued). Journal of Biological Chemistry 283, 34880–34886, doi:10.1074/jbc.M804183200 (2008).

18 Zhang, P. et al. Nontelomeric splice variant of telomere repeat-binding factor 2 maintains neuronal traits by sequestering repressor element 1-silencing transcription factor. Proc Natl Acad Sci U S A 108, 16434–16439, doi:10.1073/pnas.1106906108 (2011).

19 Liang, H., Fekete, D. M. & Andrisani, O. M. CtBP2 downregulation during neural crest specification induces expression of Mitf and REST, resulting in melanocyte differentiation and sympathoadrenal lineage suppression. Molecular and cellular biology 31, 955–970, doi:10.1128/mcb.01062-10 (2011).

20 Shimojo, M. Characterization of the nuclear targeting signal of REST/NRSF. Neuroscience Letters 398, 161–166, doi:10.1016/j.neulet.2005.12.080 (2006).

21 Shimojo, M., Lee, J. H. & Hersh, L. B. Role of zinc finger domains of the transcription factor neuron-restrictive silencer factor/repressor element-1 silencing transcription factor in DNA binding and nuclear localization. Journal of Biological Chemistry 276, 13121–13126, doi:10.1074/jbc.M011193200 (2001).

22 Chen, G.-L., Ma, Q., Goswami, D., Shang, J. & Miller, G. M. Modulation of nuclear REST by alternative splicing: a potential therapeutic target for Huntington’s disease. Journal of Cellular and Molecular Medicine, n/a-n/a, doi:10.1111/jcmm.13209.

23 Yu, M. et al. Alteration of NRSF expression exacerbating 1-methyl-4-phenyl-pyridinium ion-induced cell death of SH-SY5Y cells. Neuroscience Research 65, 236–244 (2009).

24 T, L. et al. Addendum: REST and stress resistance in ageing and Alzheimer’s disease. Nature 540,470–470, doi:10.1038/nature20579 (2016).

